# Ancestral protein reconstruction reveals the mechanism of substrate specificity in FN3K-mediated deglycation

**DOI:** 10.1101/2025.07.30.667714

**Authors:** Jenet K. Matlack, Robert E. Miner, Jameela Lokhandwala, Jennifer M. Binning

## Abstract

Protein glycation is a detrimental byproduct of living cells’ reliance on carbohydrate metabolism, and nearly all organisms encode kinases that facilitate the removal of early glycation products. In humans, these repair functions are performed by Fructosamine-3 kinase (FN3K) and Ketosamine-3 kinase (KT3K) enzymes which share conserved catalytic mechanisms but differ in substrate specificity. Recent structural studies defined key active site residues required for FN3K activity on a model substrate, yet the molecular basis for differential substrate recognition by FN3K and KT3K remains unresolved. Here, we integrate phylogenetic analysis, ancestral protein reconstruction (APR), and mutational biochemistry to to elucidate how substrate specificity evolved within the fructosamine-3 kinase family. We show that conserved substrate-binding residues are required for the phosphorylation of both fructosamines and ketosamines, but do not contribute to substrate specificity. Using APR, we resurrected four ancestral fructosamine kinases that recapitulate the distinct substrate preferences of FN3K and KT3K despite differing by only 12 amino acids. Through mutational studies and structural analysis, we show that substrate specificity is modulated by an evolutionarily tuned allosteric network that enables long-range intramolecular communication. These insights provide a new mechanistic framework for understanding FN3K substrate selection and open avenues for rational design of FN3K-selective therapeutics targeting protein glycation in metabolic disease and aging.

## Introduction

Protein glycation is a non-enzymatic post-translational modification that is implicated in many metabolic and aging-related disorders, such as diabetes, cardiovascular disease, neurodegenerative disease, and cancer^1–6^. Protein glycation occurs when reducing sugars, including glucose and ribose, react with free amine groups on amino acids to form an intermediate Schiff base. These Schiff bases can rearrange to form early glycation products called ketosamines. If left unchecked, glycation can undergo further rearrangements into advanced glycation end products (AGEs). Through receptors for advanced glycation end products (RAGEs), AGEs are activators of the NF-KB inflammatory signaling pathway, promoting both cellular damage and apoptosis^7^. While AGEs are thought to be irreversible products^8,9^, early glycation products can be enzymatically repaired via fructosamine-3 kinases (FN3Ks).

FN3Ks are conserved across the tree of life and are present in both eukaryotic and prokaryotic organisms^10^. FN3Ks repair glycated proteins by phosphorylating the O3’ of the sugar moiety on glycated amines. This generates an unstable product that reverts to the Schiff base intermediate, ultimately generating the unmodified amine, an inorganic phosphate, and byproducts such as 3-deoxyglucosone. There are two paralogues of FN3Ks in higher orders of life: FN3K and ketosamine-3 kinase (KT3K or FN3K-RP). In humans, these enzymes share 64% sequence identity at the amino acid level^11,12^. Despite this similarity, KT3Ks and FN3Ks recognize different glycated substrates. While both enzymes share overlapping activity on ribulosamines, only FN3K is capable of phosphorylating fructosamines^13,14^. This diversification of FN3K substrates later in evolution highlights the evolutionary necessity of fructosamine repair in higher orders of life^14–16^

Human FN3K (HsFN3K) has recently emerged as a therapeutic target due to its role in deglycating Nuclear factor erythroid 2–related factor 2 (NRF2), a master regulator of oxidative stress. Under normal physiologically conditions, NRF2 is tightly regulated by Kelch-like ECH-associated protein 1 (KEAP1) and acts as a transcription factor in the presence of reactive oxygen species. In many cancers, NRF2 is constitutively activated, leading to metabolic reprogramming, increased proliferation, and resistance to therapy^15,16^. Depletion of FN3K reduced tumor burden in NRF2-driven hepatocellular carcinoma and lung non-small cell lung cancer xenograft mouse models, implicating FN3K as a cellular regulator of NRF2 function^17^. Additionally, HsFN3K can directly phosphorylate ribose-glycated NRF2 peptide, further supporting NRF2 as a physiological substrate for FN3K^18^. Despite FN3K’s emerging relevance in cancer biology, the extent to which FN3K and KT3K have redundant or distinct physiological roles remains unclear. This presents a critical challenge, as it is unknown whether dual inhibition of both KT3K and FN3K would be detrimental to viability, or if selective targeting may enable a therapeutic window for FN3K antagonists. Thus, a deeper understanding of the structural and evolutionary mechanisms underlying FN3K and KT3K substrate specificity is essential for guiding rational drug design.

Recently, we and others reported the structural basis for deglycation by HsFN3K^18,19^. These studies defined a set of conserved substrate-binding residues essential for catalysis on 1-deoxy-1-morpholino-D-fructose (DMF), a small molecule fructosamine mimic. While these studies advance our understanding of FN3K protein deglycation, they did not address the distinct yet overlapping substrate specificities of FN3K and KT3K. The structural and evolutionary basis for this functional divergence remains unresolved. In this study, we address this gap by integrating phylogenetic analysis, ancestral protein reconstruction (APR), and mutational biochemistry to elucidate the molecular determinants of substrate specificity within the fructosamine-3 kinase family. We show that substrate binding residues in HsFN3K are conserved across all fructosamine-3 kinases and are required for the phosphorylation of both fructosamines and ketosamines. Using APR, we reconstructed ancestral enzymes that recapitulate the substrate preferences of HsFN3K and HsKT3K, despite differing by only 12 amino acids. Through systematic mutagenesis, we identified that substrate specificity in FN3Ks is regulated not by direct interactions within the active site, but by an allosteric network that governs fructosamine recognition through long-range intramolecular communication. By defining how FN3Ks evolved to selectively recognize glucose-glycated proteins, we lay the foundation for structure-guided therapeutic targeting of this clinically relevant enzyme.

## Results

### Identification of FN3K residues required for glycated protein recognition

Recent high-resolution structural studies have identified key residues within HsFN3K that mediate binding to the substrate mimetic DMF (Figure 1A). Site-directed mutagenesis confirmed the functional importance of W219, H288, and H291 in binding DMF and substrate recognition^18,19^. To identify additional residues contributing to substrate engagement across the FN3K family, we performed a ligand-protein interaction analysis using the HsFN3K–ATP–DMF pre-catalytic complex structure. This analysis revealed nine DMF-contacting residues within the substrate-binding pocket that contact DMF: I25, D217, W219, F252, N287, H288, H291, F292, and G152 (Figure 1B). Among these residues, N287 stabilizes the sugar moiety through hydrogen bonding, allowing D217 to function as the catalytic base. The side chains of W219, F252, H288, H291, and F292 form stabilizing interactions with the substrate, while G152 contributes through backbone interactions.

**Figure 1.**
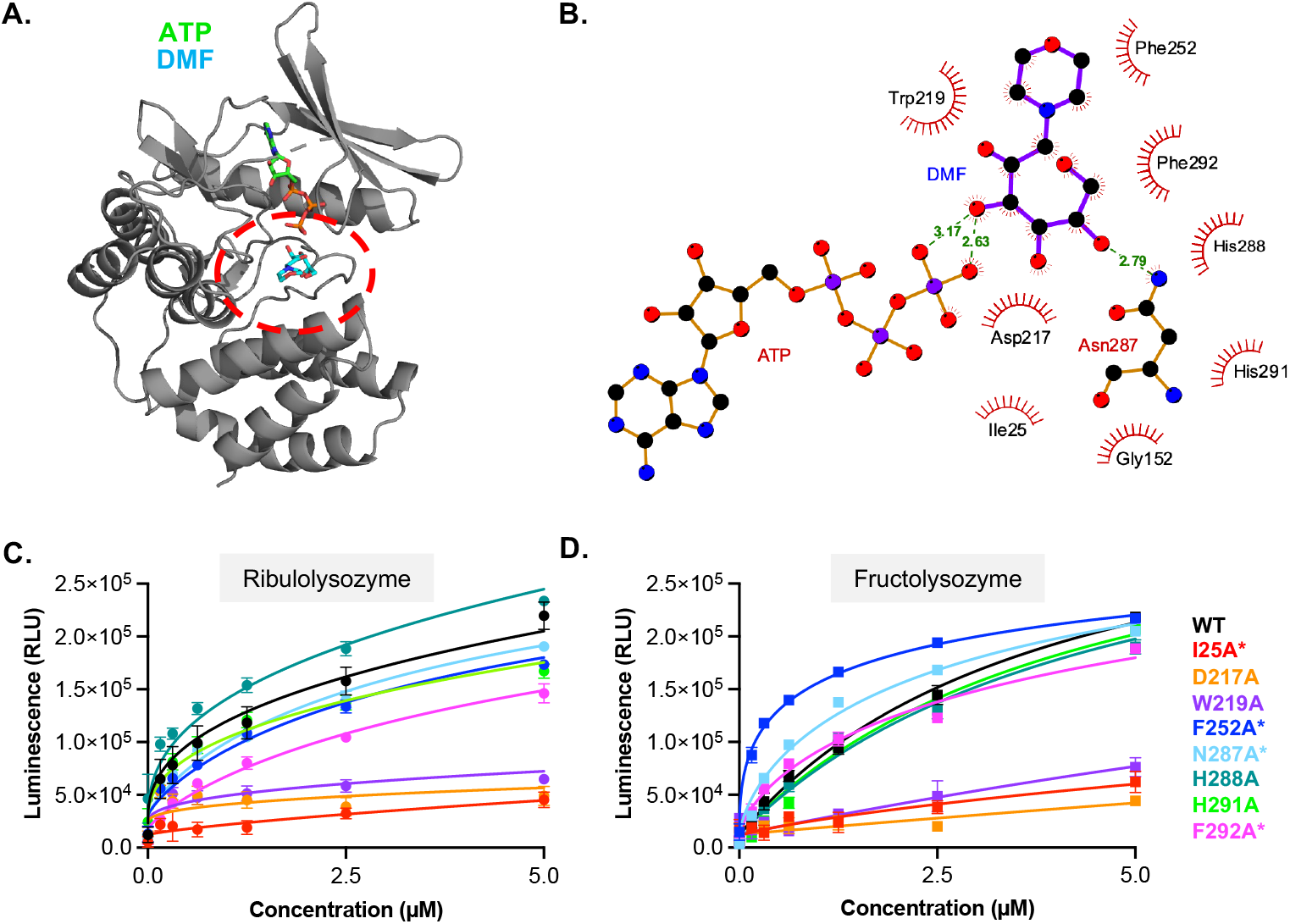
Substrate binding residues in HsFN3K are required for small molecule and protein substrates. **A**. Ribbon diagram of HsFN3K bound to DMF and ATP (PDB: 9CXM). The substrate binding pocket is highlighted in red. **B**. Ligand-protein interaction analysis of HsFN3K with DMF and ATP. **C-D**. Kinase assay of HsFN3K WT, I25A, D217A, W219A, F252A, N287A, H288A, H291A, and F292A on ribulolysozyme and fructolysozyme. HsFN3K was titrated in the indicated concentration range and ATP consumption was measured by luminescence. Each data point represents means of triplicates; error bars indicate standard error. Asterisks (^*^) indicates that the protein has an MBP-tag.

To evaluate the functional relevance of these contacts, each residue was individually mutated to alanine, and kinase activity was assessed using a luciferase-based assay. Unlike previous studies, which primarily utilized DMF, we used fructosamine- and ribulosamine-modified lysozyme to assess FN3K activity against glycated proteins (Figure 1C-D). We observed that only I25A, D217A, and W219A lost activity towards both fructo- and ribulolysozymes. We previously showed that D217A and W219A retain ATP binding but fail to interact with DMF^19^. Although I25 is solvent-exposed in the apo structure, it reorients to the catalytic pocket in the ternary complex, such that it is positioned at the interface of between ATP and DMF (Supplemental Figure 1D). Using differential scanning fluorimetry (DSF), we qualitatively assessed the ability of HsFN3K I25A to bind ATP (Supplemental Figure 1E). The I25A mutant fails to exhibit ATP-induced thermal stabilization, suggesting a critical role for I25 in positioning both ATP and substrate for catalysis.

Our data also established that HsFN3K F252A, N287A, H288A, H291A, and F292A mutants retain activity towards both glycated lysozymes. Consistent with our previous study, which showed that HsFN3K H288A impaired but did not abolish kinase activity towards DMF, this mutation showed a slight decrease in activity towards fructolysozyme. Interestingly, this mutation showed an increase in activity towards ribulolysozyme. Similarly, we observed F252A and N287A mutants have opposing effects on the catalytic activity of HsFN3K towards the two glycated substrates, with these mutations enhancing activity in the presence of fructosamines, but not ribulosamines.

Together, these data define the HsFN3K residues critical for substrate binding and HsFN3K activity. Of the nine residues that contact DMF, only I25, D217, and W219 are essential for catalytic activity, suggesting that these residues function as universal substrate recognition elements within the FN3K catalytic pocket.

### Conserved substrate-binding residues do not account for differences in substrate specificity between FN3Ks and KT3Ks

Previous phylogenetic and biochemical analyses have shown that both mammalian and non-mammalian FN3Ks can phosphorylate fructosamines, whereas KT3Ks display a narrower substrate preference (Figure 2A)^13,14^. Conservation analysis across >14,000 fructosamine-3 kinase family members revealed that all nine substrate-binding residues identified in the HsFN3K-DMF co-structure are highly conserved across the family (Figure 2B). This conservation persists in both FN3Ks and KT3Ks, despite their overlapping yet distinct substrate specificities. We next compared global sequence identities within and between the two subfamilies. While FN3Ks and KT3Ks exhibit high sequence conservation within each lineage, percent identity drops significantly when comparing members across the two clades (Figure 2C).

**Figure 2.**
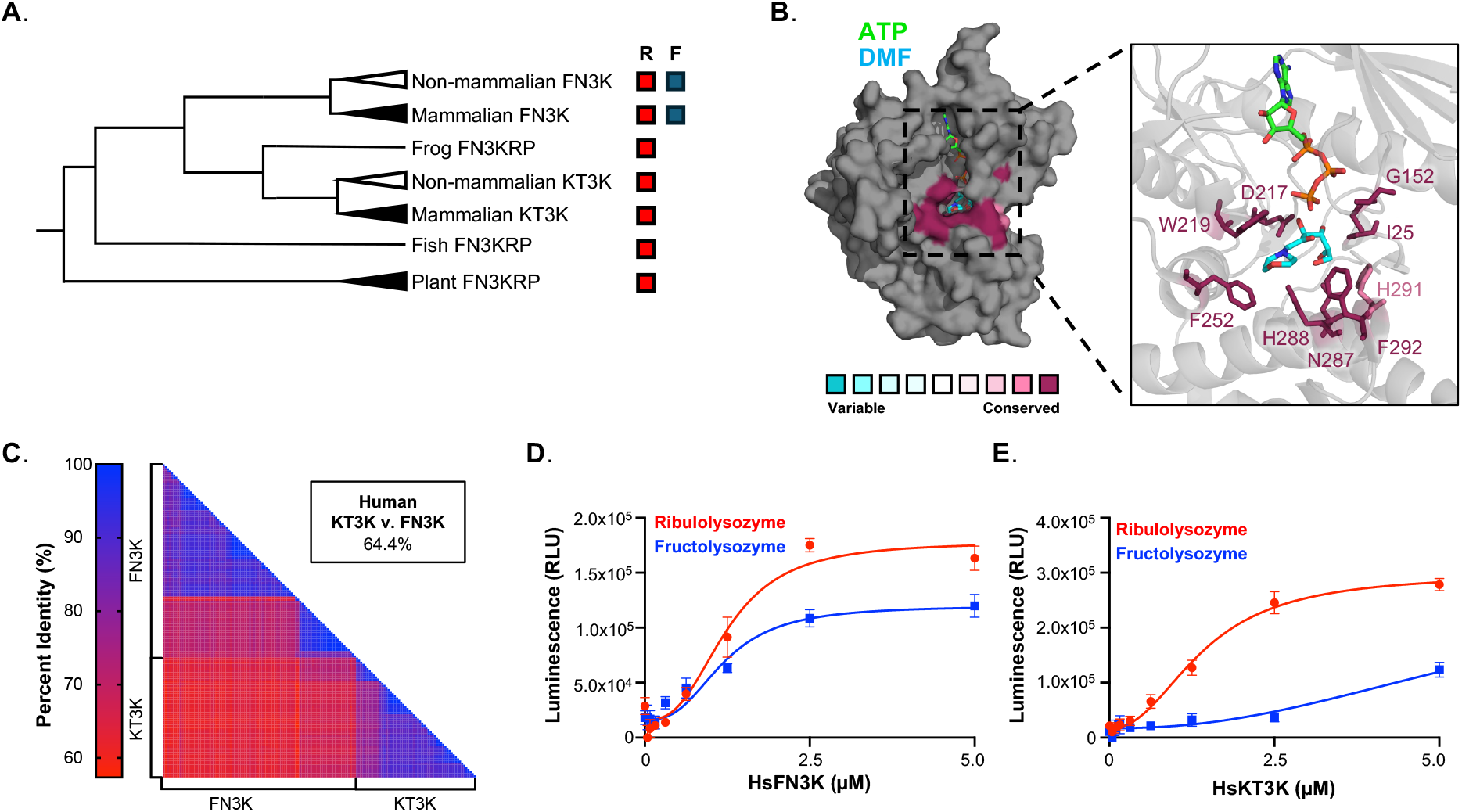
Substrate specificity differs between FN3K and KT3K despite conservation of substrate binding residues. **A**. Unrooted phylogeny tree of fructosamine-3 kinase family. Substrate specificity towards ribulosamine (R) and fructosamine (F) is indicated on the right of the phylogeny tree. **B**. Conservation analysis of fructosamine kinase family represented on HsFN3K bound to DMF and ATP (PDB: 9CXM). The colors indicate level of conservation of residues according to the scale. Substrate binding residues are highlighted in the inlet. **C**. Percent identity matrix of amino acid sequences of FN3Ks and KT3Ks post-duplication. Colors are represented by the scale bar. **D-E**. Kinase assay of HsFN3K (D) and HsKT3K (E) on ribulolysozyme (red) and fructolysozyme (blue). Enzyme was titrated in the indicated concentration range and ATP consumption was measured by luminescence. Each data point represents means of triplicates; error bars indicate standard error.

To experimentally validate differences in substrate preference, we expressed and purified recombinant HsFN3K and HsKT3K and tested their activity using a luciferase-based kinase assay. As expected, HsFN3K phosphorylated fructolysozyme (Figure 2D), whereas HsKT3K exhibited no detectable activity toward this substrate (Figure 2E). Both enzymes retained activity on ribulolysozyme, a common fructosamine-3 kinase substrate. These data demonstrate that the amino acids directly involved in substrate binding are conserved across the fructosamine-3 kinase family, and therefore do not account for the divergent substrate specificities observed between FN3Ks and KT3Ks. These findings suggest that additional residues, likely acting outside the substrate-binding pocket, underlie the evolutionary diversification in substrate preference.

### Ancestral enzymes recapitulate origins of FN3K substrate specificity

To identify the evolutionary adaptations underlying FN3K and KT3K substrate specificity, we employed ancestral protein reconstruction (APR) to resurrect ancestral fructosamine kinases that recapitulate the substrate preferences of human FN3K and KT3K. This approach integrates information from multiple sequence alignments and phylogenetic trees to infer the most probable ancestral sequences at key evolutionary nodes of divergence in the tree^20,21^. Given that FN3K emerged after fish speciation, sequences from bacterial orthologues were excluded, and phylogenetic analysis began with plant orthologues. From this, we identified four ancestral proteins of interest (Figure 3A): ancFN3KRP, which predates the gene duplication event; ancKT3K^1^ and ancKT3K^2^, both from the KT3K lineage; and ancFN3K, which lies on the FN3K lineage preceding the emergence of HsFN3K. All four ancestral sequences preserved the nine substrate-contacting residues identified in Figure 1B (Supplemental Figure 2A).

**Figure 3.**
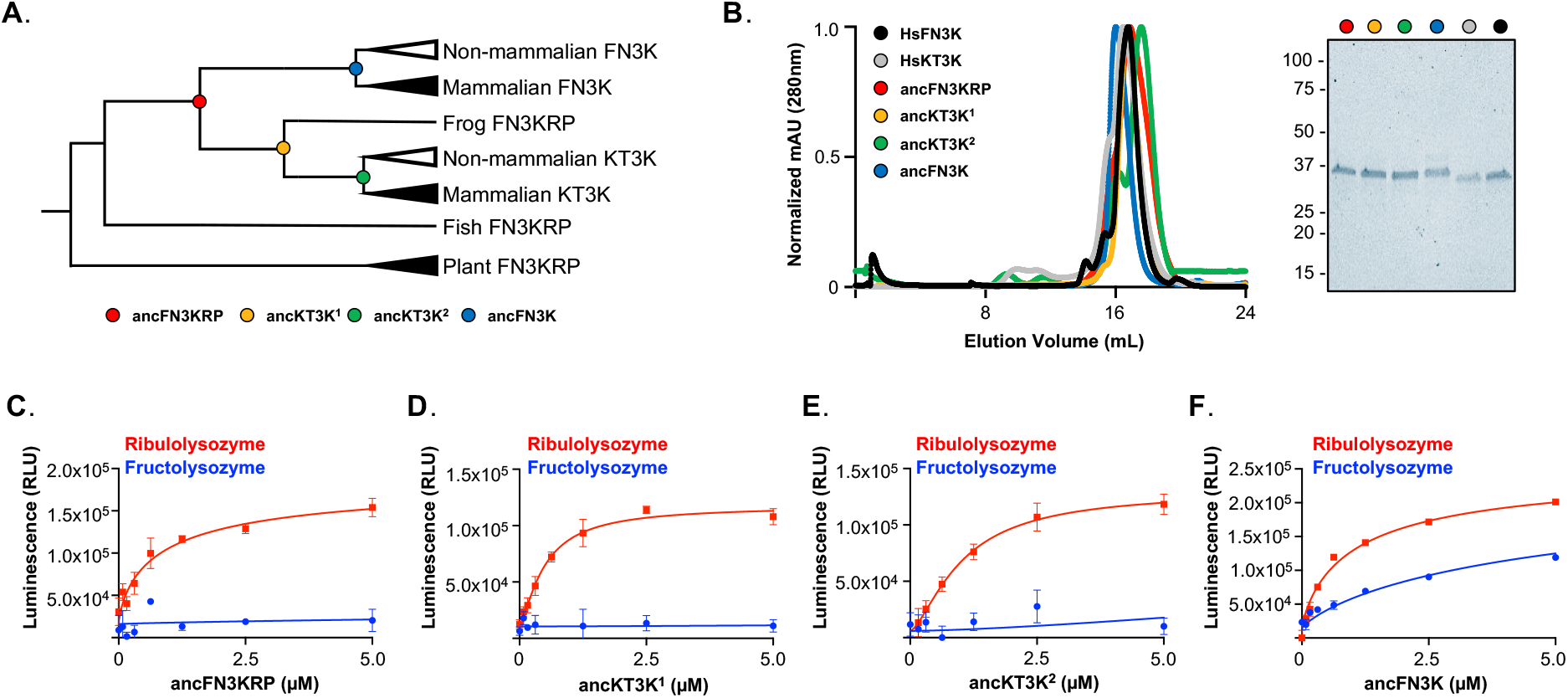
Ancestral reconstruction of fructosamine-3 kinase family recapitulates substrate specificity. **A**. Unrooted phylogeny tree of fructosamine-3 kinase family with locations of identified ancestral proteins. ancFN3KRP (red), ancKT3K^1^ (yellow), ancKT3K^2^ (green), and ancFN3K (blue) are represented as colored circles on the tree. **B**. Size-exclusion chromatography profile of HsFN3K, HsKT3K, ancFN3KRP, ancKT3K^1^, ancKT3K^2^, and ancFN3K. The inset shows the Coomassie stained SDS-PAGE of each protein. **C-F**. Kinase assay of ancFN3KRP (C), ancKT3K^1^ (D), ancKT3K^2^ (E), and ancFN3K (F) on ribulolysozyme (red) and fructolysozyme (blue). Enzyme was titrated in the indicated concentration range and ATP consumption was measured by luminescence. Each data point represents means of triplicates; error bars indicate standard error.

Each ancestral protein was recombinantly expressed and purified, eluting as a monodisperse peak by size-exclusion chromatography at a similar elution volume to HsFN3K and HsKT3K (Figure 3B). Thermal stability analysis revealed that ancKT3K^1^ and ancKT3K^2^ displayed equivalent or increased melting temperatures compared to HsKT3K. This is consistent with the well-documented “consensus effect” in APR, whereby ancestral sequences tend to exhibit enhanced solubility and stability due to the enrichment of high-probability amino acids within the protein family^22^. This is not always the case, however, as ancFN3K had a markedly lower melting temperature compared to HsFN3K (Supplemental Figure 2B-C).

We next assessed the catalytic activity of these reconstructed proteins using glycated lysozymes as substrates. All four ancestral proteins retained activity towards ribulolysozyme, but only ancFN3K exhibited activity towards fructolysozyme (Figure 3C-F). These results demonstrate that the reconstructed ancestral enzymes recapitulate substrate preferences observed in extant KT3Ks and FN3Ks, and that the ability to repair fructosamines may have emerged specifically within the FN3K lineage.

### Reconstructing the neofunctionalization of fructosamine phosphorylation in FN3Ks

To identify the molecular determinants that underlie the substrate specificity differences between FN3Ks and KT3Ks, we focused on two reconstructed ancestral enzymes: ancFN3KRP and ancFN3K. These two ancestral proteins recapitulate the substrate preferences observed in their extant counterparts despite differing at only 12 amino acid positions (Figure 4A). All 12 residues are located on the periphery of the substrate-binding pocket (Figure 4B), providing a focused framework to dissect the molecular basis of substrate specificity between FN3K and KT3K.

**Figure 4.**
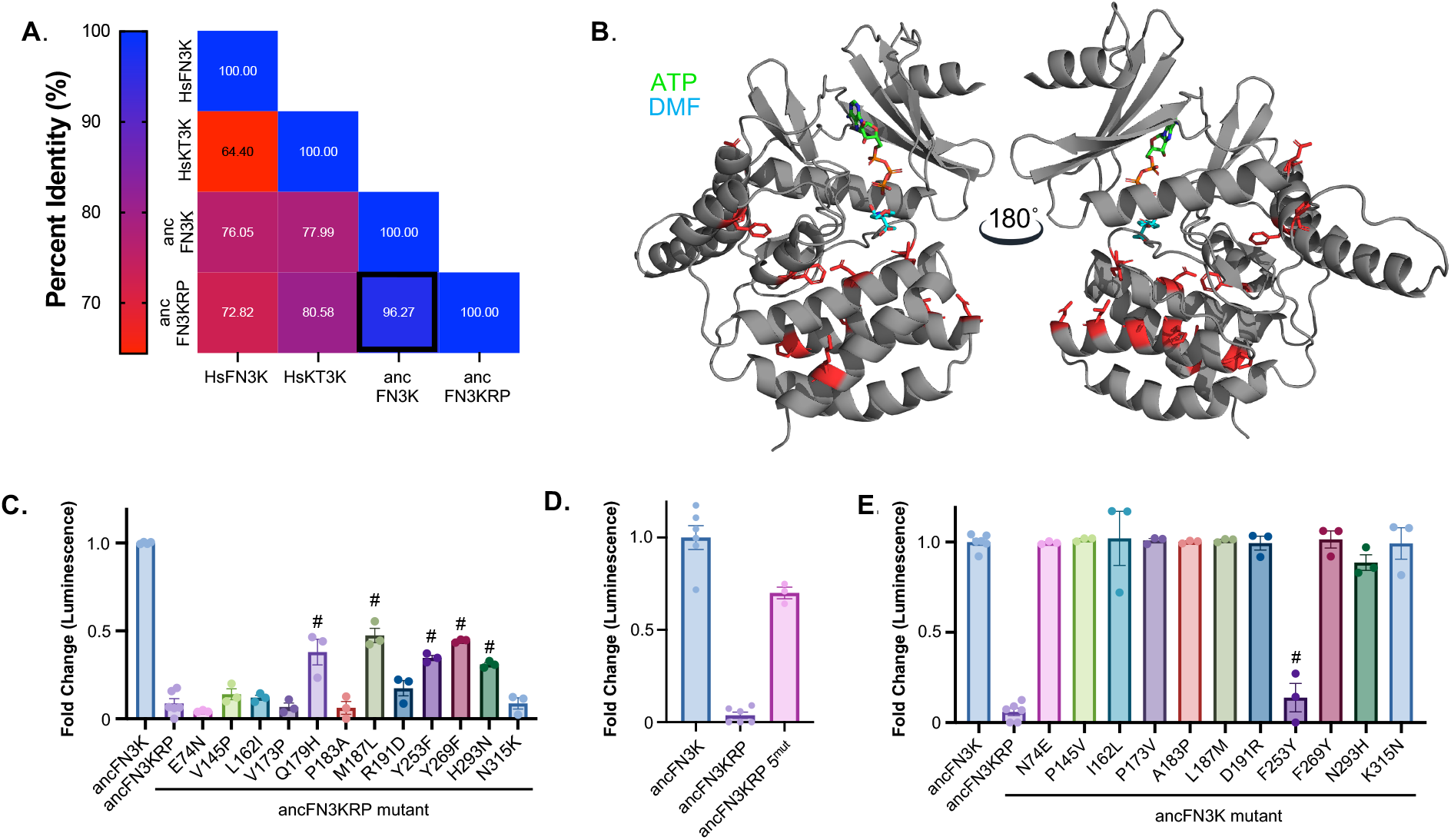
Systematic mutagenesis of ancFN3KRP and ancFN3K. **A**. Percent identity matrix of HsFN3K, HsKT3K, ancFN3KRP, and ancFN3K **B**. AlphaFold3 model of ancFN3K. DMF and ATP were modeled into the structure by aligning HsFN3K ATP-DMF with ancFN3K in PyMOL. The 12 amino acid residues that differ between ancFN3K and ancFN3KRP are highlighted in red as sticks. **C-E**. Kinase assay of ancFN3KRP GOF (C and (D) and ancFN3K LOF (E) mutations against DMF. Each bar graph represents the activity at 5 µM of enzyme. Each data point represents means of triplicates; error bars indicate standard error. P-value was determined using Dunnett’s multiple comparisons test, #: p-value < 0.0001.

To elucidate the evolutionary changes that enabled deglycation of fructosamines, we conducted reciprocal mutagenesis to functionally convert ancFN3KRP into ancFN3K (gain-of-function) and ancFN3K into ancFN3KRP (loss-of-function) by introducing the 12 residue substitutions individually and in combination. We assessed kinase activity using the model substrate DMF, which enables quantitative measurement of FN3K-like activity. Among the 12 amino acid substitutions, five mutants (Q179H, M187L, Y253F, Y269F, and H293N) conferred enzymatic activity toward DMF when introduced into ancFN3KRP, indicating partial gain of FN3K-like specificity (Figure 4C). While none of the single substitutions restored activity to the level of wild-type ancFN3K, combining all five substitutions (ancFN3KRP^-^5^mut^) further enhanced activity toward DMF, reaching approximately 70% of ancFN3K wild-type levels (Figure 4D). Importantly, activity toward ribulolysozyme was retained for all gain-of-function mutations, establishing that these mutations do not broadly disrupt catalytic activity (Supplemental Figure 3B-C).

We next performed complementary loss-of-function mutagenesis in the ancFN3K background to identify substitutions that would reduce or eliminate fructosamine activity, thereby rendering the enzyme more KT3K-like. Each of the 12 residues were individually reverted to their ancFN3KRP identity. In contrast to the gain-of-function results, only one substitution, F253Y, resulted in a significant reduction in DMF activity compared to wild-type ancFN3K (Figure 4E), suggesting that F253 plays a central role in maintaining fructosamine specificity. Notably, the H179Q variant was insoluble in *E. coli* expression systems and could not be purified for activity assays. Together, these data demonstrate that substrate specificity divergence between FN3K and KT3K arose from a small number of amino acid substitutions following gene duplication. Among these, F253 (corresponding to HsFN3K F244) emerged as a key determinant in fructosamine recognition in the FN3K lineage.

### Substrate specificity in HsFN3K is modulated by peripheral residues identified in ancestral enzymes

To investigate whether residues identified through ancestral protein reconstruction contribute to substrate specificity in HsFN3K, we first examined their conservation across the FN3K family. All five residues are highly conserved among both mammalian and non-mammalian FN3Ks and KT3Ks (Figure 5A). Moreover, the residues surrounding these are also well-conserved, suggesting that substrate recognition is governed by a tightly constrained structural environment.

**Figure 5.**
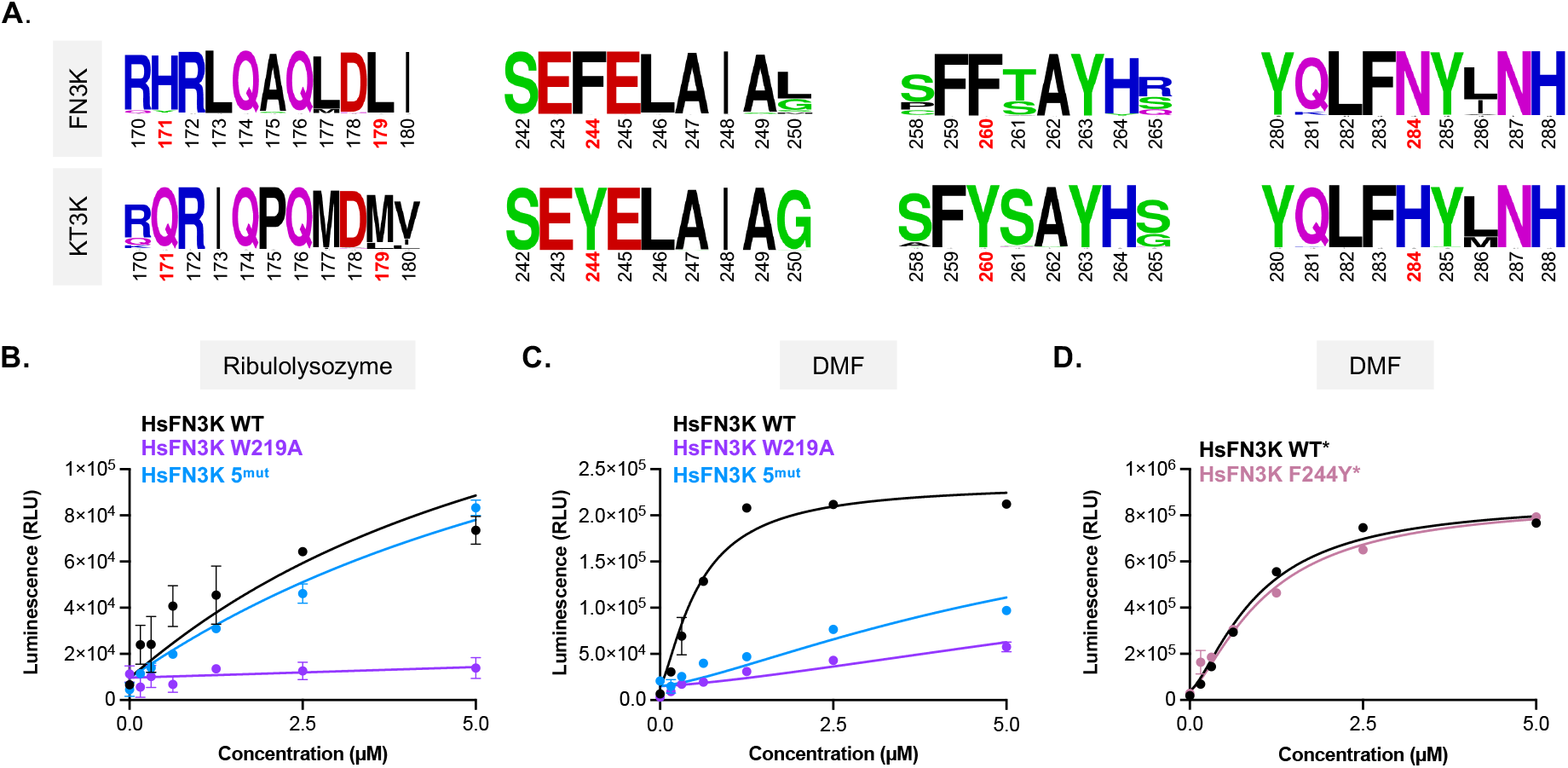
Identified residues in ancestral proteins cause loss of function in human FN3K. **A**. LogoPlot of FN3Ks and KT3Ks. HsFN3K 5^mut^ amino acid residues identified in ancestral proteins are highlighted in red. **B-C**. Kinase assay of HsFN3K WT, W219A, and 5^mut^ against ribulolysozyme (C) and DMF (C). **D**. Kinase assay of HsFN3K WT^*^ and HsFN3K F244Y^*^. Asterisks (^*^) indicates that the protein has an MBP-tag. Each data point represents means of triplicates; error bars indicate standard error.

To determine whether these residues regulate substrate specificity in HsFN3K, we introduced the corresponding ancestral substitutions into HsFN3K to generate a five-point mutant, HsFN3K 5^mut^ (H171Q/M179L/F244Y/F260Y/N284H). We then assessed enzymatic activity against both ribulolysozyme and DMF. HsFN3K 5^mut^ retained activity towards ribulolysozyme (Figure 5B), but lost activity toward DMF (Figure 5C). The combination of five mutations is sufficient to restrict activity to ribulosamines, recapitulating the narrower substrate preference characteristic of KT3K. Additionally, HsFN3K 5^mut^ exhibited an 8°C increase in melting temperature relative to wild-type HsFN3K (Supplemental Figure 4), consistent with a more conformationally stable, KT3K-like state. Given that a single mutation (F253Y) in the ancestral enzyme was sufficient to restrict DMF activity, we next tested whether the analogous mutation in HsFN3K (F244Y) could independently recapitulate this effect. In contrast to the ancestral context, HsFN3K F244Y retained activity toward DMF (Figure 5D), suggesting establishing that this mutation alone is insufficient to alter substrate specificity in the human enzyme. Together, these data demonstrate that multiple adaptations on the periphery of the substrate binding pocket contribute collectively to substrate specificity in HsFN3K. These findings underscore the importance of cooperative interactions between distal residues in shaping functional divergence within the FN3K family.

### Intramolecular interaction analysis reveals dynamic allosteric regulation through residue 244 in FN3Ks and KT3Ks

Structural mapping of the HsFN3K 5^mut^ residues onto the ATP-DMF-bound HsFN3K structure revealed that none make direct contact with the substrate or substrate-binding residues (Figure 6A). This suggests that these amino acids influence activity through indirect mechanisms, potentially via allosteric regulation of the active site. Comparison of the apo and substrate-bound HsFN3K structures showed no substantial conformational changes upon ligand binding (Supplemental Figure 5A), indicating that the determinants of specificity may be pre-configured in the apo state.

**Figure 6.**
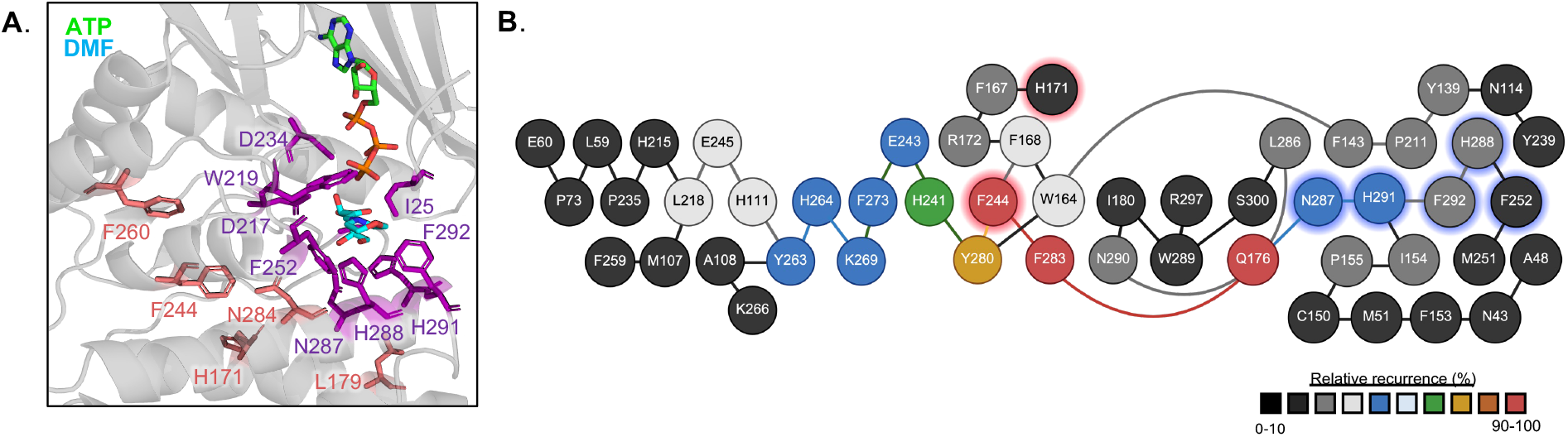
Protein Structure Network analysis of HsFN3K. **A**. Substrate binding pocket of HsFN3K ATP-DMF structure. HsFN3K 5^mut^ residues are highlighted in salmon. DMF binding residues are highlighted in purple. **B**. WebPSN network analysis of HsFN3K (PDB: 9CX8). Red highlights signify residues identified in this paper. Blue highlights signify substrate binding residues. Relative recurrence represents the strength of interactions and is indicated by the scale bar. Node colors are representative of the recurrence in network analysis and follow the same scale.

To explore whether these residues participate in a secondary interaction network influencing the substrate-binding site, we examined residue-level interactions within a 6 Å cutoff. Residues 171, 179, 244, and 260 exhibited no contacts with substrate-binding residues in either HsFN3K or HsKT3K. Although residue 284 formed interactions with H288, N287, F252, and D217, key amino acids in the HsFN3K substrate-binding pocket, these interactions were conserved in both HsFN3K and HsKT3K and did not differ in distance or geometry, suggesting that residue 284 does not contribute to specificity through direct structural interactions (Supplemental Figure 5B).

Given the well-established role of long-range, allosteric networks in kinase regulation^23,24^, we next performed Protein Structure Network (PSN) analysis to evaluate potential allosteric signaling within FN3K (Figure 6B). Using a 3 Å distance cutoff, we identified the shortest communication pathways that pass through the HsFN3K 5^mut^ residues. HsFN3K F244 and H171 are predicted to form an interaction network that extends to five of the nine substrate-binding residues, forming a distinct communication pathway not found in KT3K-like proteins, such as A. thaliana FN3K (AtFN3K) (Supplemental Figure 6A-B). Together, these results support our mutational studies and provide a framework for how the five specificity-determining residues may function through dynamic allosteric networks rather than direct substrate engagement. In particular, residue 244 emerges as a central regulator of long-range intramolecular communication, facilitating a shift in allosteric connectivity that modulates substrate specificity in FN3Ks and KT3Ks.

## Discussion

Protein kinases represent one of the most therapeutically exploited classes of enzymes in the human proteome. FN3K has emerged as a therapeutic target due to its role in protein repair and regulation of redox signaling. Notably, FN3K has been implicated in the post-translational control of NRF2, a master regulator of oxidative stress responses and a known driver of chemoresistance in several cancers^25–28^. Given the long-standing challenges associated with developing NRF2 antagonists, there is a growing interest in identifying alternative therapeutic strategies for NRF2-driven cancers^29^. Recent studies have demonstrated that NRF2 is susceptible to non-enzymatic glycation, and that FN3K-mediated deglycation can restore its transcriptional activity. These findings highlight FN3K as a potential vulnerability in NRF2-addicted tumors. However, to fully exploit FN3K as a therapeutic target, we need a detailed understanding of the molecular mechanisms underlying FN3K’s function, particularly its substrate specificity, which to date has remained poorly defined.

In this study, we describe the molecular basis of substrate specificity in the FN3K family. Although FN3K and KT3K share overlapping activity on ribulosamines, only FN3K is capable of phosphorylating fructosamines. Using the recently published HsFN3K-ATP-DMF pre-catalytic structure, we identified nine residues that coordinate DMF, a fructosamine mimetic, within the active site of HsFN3K. We performed a conservation analysis, which revealed that all nine substrate-binding residues are highly conserved across the FN3K family, suggesting that molecular features beyond the substrate binding pocket govern substrate selectivity.

Through a combination of structural analysis, ancestral protein reconstruction (APR), and biochemical characterization, we demonstrate that substrate specificity is not regulated by FN3K active site residues, but instead by a peripheral allosteric network. By using APR, we generated four ancestral proteins that recapitulated the evolution of FN3Ks with minimal divergence in sequence. This approach allowed us to systematically identify the amino acid substitutions that enabled the acquisition of FN3K-specific activity. Despite 96% sequence identity, ancestral proteins from the pre- and post-gene FN3KRP duplication event (ancFN3KRP and ancFN3K, respectively) displayed distinct substrate preferences, mirroring what is observed in the HsFN3K and HsKT3K. Through gain- and loss-of-function mutagenesis, we identified five amino acids (Q179H, M187L, Y253F, Y269F, and H293N) that regulate substrate specificity. Remarkably, a single adaptation at position 253 (in ancFN3K/ancFN3KRP) serves as a specificity switch that governs activity towards fructosamines. However, this single adaptation in HsFN3K was not sufficient to alter substrate specificity. While the thermal stability of ancFN3KRP was comparable to that of HsFN3K, the ancFN3KRP 5^mut^, which gained the ability to deglycate fructosamines, resulted in a 16°C decrease in melting temperature. This observation is consistent with prior studies showing that the evolution of new enzymatic functions is often accompanied by a loss of protein stability^30–33^. Our findings suggest that additional stabilizing mutations likely arose during the evolutionary transition from ancFN3K to HsFN3K (T_m_ = 46°C and 56°C, respectively), compensating for destabilizing changes associated with the expanded substrate specificity toward fructosamines. These results provide insight into the evolutionary requirements for the emergence of fructosamine as a physiological substrate for FN3Ks.

To build on our findings from ancestral protein reconstruction, we investigated the functional impact of the corresponding gain- and loss-of-function mutations in HsFN3K. Since the single HsFN3K F244Y substitution was insufficient to reduce activity toward DMF, we tested the combined effect of five mutations identified in our ancestral proteins. Introducing all five substitutions into HsFN3K successfully restricted its activity to ribulosamines, thereby mimicking the narrower substrate specificity observed in KT3K. This shift indicates that substrate preference in HsFN3Ks is governed by a distributed network of residues that modulate catalysis through structural and dynamic effects, rather than by a single determinant.

Protein structure network analysis further supports this model, revealing long-range intramolecular networks with the substrate-binding residues. Specifically, in HsFN3K, residue F244 connects to F283 and propagates through Q176 to five of the nine substrate-interacting residues, suggesting an allosteric pathway. Although the F244Y mutation alone does not alter enzyme activity, both our ancestral reconstruction and PSN analyses identify F244 as a key interaction node within a broader regulatory network that modulates substrate specificity. Together, these findings support a model in which amino acids outside the active site, acting through an allosteric network, fine-tune substrate specificity in the FN3K family.

These findings have important implications for drug discovery efforts targeting FN3K. The high conservation of the active site across FN3Ks and KT3Ks suggests that ATP-competitive or substrate-mimetic inhibitors will likely target both enzymes indiscriminately. This raises the possibility that FN3K inhibition could lead to off-target effects, particularly in tissues where KT3Ks may have independent functions. Additional studies that define the biological role of FN3K and KT3K will be critical to understand the therapeutic consequences of a pan-FN3K inhibitor. Moreover, our separation-of-function mutations provide powerful tools for dissecting the physiological roles of FN3K in vivo. Future studies can now ask whether fructosamine or ribulosamine repair is more critical in specific tissues or disease states. This will be especially valuable in models of metabolic dysfunction or NRF2-driven cancers. Ultimately, understanding how FN3K’s substrate preferences are wired into its structure opens new avenues for therapeutic intervention and illuminates the broader principles of kinase substrate specificity.

## Methods

### Plasmids

HsFN3K WT, D217A, W219A, H288A, and H291 were cloned previously^19^. All constructs were made using isothermal assembly and standard PCR techniques. Ancestral proteins were codon-optimized for *E. coli* expression (Integrated DNA Technologies) and cloned into a modified pET15b vector containing a N-terminal maltose-binding protein and His-tag. HsKT3K was cloned into the pFastBac™ Dual vector (Gibco #10712024) with mCherry in the second multiple cloning site. pGro7 was obtained from Takara Bio (#3340).

### Protein expression and purification

HsFN3K and ancestral proteins were co-expressed with the pGro7 plasmid containing GroEL/GroES in BL21(DE3) *E. coli* cells. Cells were grown to an OD of 0.5 - 0.7 and induced with 0.5 mM IPTG and 0.5 mg/mL L-arabinose for 16 hours at 18°C. Cells were harvested at 6,000 x *g*, lysed, and clarified at 40,000 x *g* in lysis buffer containing 20 mM Tris pH 8.0, 500 mM NaCl, 20 mM imidazole, 1 mM PMSF, 1 mM DTT. The clarified lysate was then subjected to Ni-NTA (Cytiva #17525501) affinity purification and eluted with a linear gradient of imidazole (20 mM – 500 mM imidazole in 50 mM Tris pH 8.0, 300 mM NaCl, 5 mM BME). Subsequently, MBP-tagged proteins were purified using MBPTrap HP (Cytiva #28918779) and eluted using 50 mM Tris pH 8.0, 300 mM NaCl, 1% maltose, and 5 mM BME. Unless otherwise stated, MBP-tags were cleaved using rTEV protease. The fusion tag was removed through Ni-NTA purification and cleaved protein was subjected to SD200 size-exclusion chromatography in a final buffer of 50 mM HEPES pH 7.0, 250 mM NaCl, and 1 mM DTT.

KT3K-His was expressed in Sf9 insect cells. Cells were harvested at a viability between 30-50%. Cell pellets were washed with ice-cold PBS supplemented with protease inhibitors (Thermo Scientific #A32961) to remove EDTA. Protein was extracted through homogenization in lysis buffer and subsequent centrifugation. Soluble fractions were purified on an AKTA using Ni-NTA affinity chromatography and SD200 size-exclusion chromatography. All proteins were concentrated to ∼4 mg/mL and flash frozen in liquid nitrogen. Proteins were stored at -80°C.

### Lysozyme glycation and colorimetric assay

60 mg/mL lysozyme (Hampton Research #HR7-110) was incubated with 4 M ribose or glucose in deionized water at 50 °C for 72 hours or 28 days, respectively. Glycation reactions were quenched through overnight dialysis against water to remove excess free sugars. Glycation was evaluated through a nitro-blue tetrazolium (NBT) assay, as previously described^34,35^. Briefly, 50 µM of substrate was incubated with 0.25 mg/ml NBT (Fisher Scientific #N6495) for 30 minutes at 37°C. The incubated mixture was diluted with 0.1 M Carbonate/Bicarbonate pH 10.3 (Thomas Scientific #C988L92) to a final concentration of 5 µM of substrate and incubated for an additional 30 minutes at 37°C. Reactions were measured at 530 nm using BioTek Synergy H1 microplate reader (Agilent Technologies). All experiments were performed in triplicate and the data were plotted using GraphPad Prism.

### Luciferase-based kinase assay

The kinase reaction was carried out at room temperature in 10 µL containing 200 µM lysozyme (lysozyme, ribulolysozyme, or fructolysozyme) or 100 µM DMF, 100 µM ATP, 5 mM MgCl2, 0.1 µM BSA, and 0-5 µM of serially diluted kinase. The reaction was quenched after 30 minutes using the Kinase-Glo® Plus reagent (Promega #V3771). The final reaction was incubated for 10 minutes at room temperature. Luminescence was measured using a BioTek Synergy H1 microplate reader (Agilent Technologies). All experiments were performed in triplicate, and the data were plotted using GraphPad Prism. Due to the possible background activity, unmodified lysozyme was used as a background reading and removed.

### Differential Scanning Fluorimetry

Differential scanning fluorimetry was performed as described previously^19^. Briefly, protein was diluted in 50 mM HEPES pH 7.0, 250 mM NaCl, 1 mM DTT, and 2 mM MgCl2 to a final concentration of 5 µM. ATP was added to a final concentration of 100 µM. 5x SYPRO dye was added before being analyzed using a StepOnePlus Real-Time PCR machine (Applied Biosystems). The mixture was heated from 20°C to 95°C at a rate of 1°C per minute. Melting curves were generated from the first derivative of the fluorescent readings. DSF scans of all samples were performed in triplicate and the data were plotted using GraphPad Prism.

### Ancestral protein reconstruction

Protein sequences for FN3Ks and KT3Ks were retrieved from UniProt^36^. Homologs were identified using the HMMER web server^37^. Multiple sequence alignments were performed using **MU**ltiple **S**equence **C**omparison by **L**og-**E**xpectation (MUSCLE)^38,39^. Phylogeny tree was generated using PhyML 3.0 using the substitution model Q.plant +G+I as selected by smart model selection (SMS) using Akaike Information Criterion (AIC)^40,41^. Ancestral sequences were predicted using Phylogenetic Analysis by Maximum Likelihood (PAML) 4.10.1 using the default settings^42^. The resulting phylogeny tree was visualized using FigTree v.1.4.4 to evaluate the nodes of interest.

### Percent Identity Matrix

Protein sequences of FN3Ks and KT3Ks were aligned using Clustal Omega^43^. Logo plots were generated using this sequence alignment in WebLogo 3.0^44^. The percent identity output was graphed using GraphPad Prism

### Protein structure and network predictions

The structures of ancFN3K and HsKT3K were predicted using AlphaFold3^45^. Protein Structure Network analyses were done using webPSN^46^. Intramolecular forces were assessed using ProteinTools^47^.

## Acknowledgments

We thank members of the Binning and Luca lab for their suggestions and technical assistance. The authors also acknowledge the Chemical Biology Core at the H. Lee Moffitt Cancer Center & Research Institute, which is supported in part by a National Cancer Institute (NCI) support grant (P30-CA076292). The protein structure network was adapted using Biorender.com. This study was supported by the U.S. National Institutes of Health grant R35 GM143004 to J.M.B.

## Author contributions

J.K.M. performed all structural and biochemical studies of FN3Ks and variants, contributed to the experimental design, and wrote the manuscript. R.E.M. performed the colorimetric assay on glycated substrates. J.L. assisted with experimental design and manuscript preparation. J.M.B. conceptualized, supervised, provided resources, reviewed, and edited the manuscript. All authors read and approved the final manuscript.

## References

1. Ahmed, M. U., Thorpe, S. R. & Baynes, J. W. Identification of N epsilon-carboxymethyllysine as a degradation product of fructoselysine in glycated protein. Journal of Biological Chemistry 261, 4889–4894 (1986).

2. Teerlink, T., Barto, R., Ten Brink, H. J. & Schalkwijk, C. G. Measurement of Nε-(Carboxymethyl)lysine and Nε-(Carboxyethyl)lysine in Human Plasma Protein by Stable-Isotope-Dilution Tandem Mass Spectrometry. Clin Chem 50, 1222–1228 (2004).

3. Krook, M., Ghosh, D., Strömberg, R., Carlquist, M. & Jörnvall, H. Carboxyethyllysine in a protein: native carbonyl reductase/NADP(+)-dependent prostaglandin dehydrogenase. Proceedings of the National Academy of Sciences 90, 502–506 (1993).

4. Sell, D. R., Lapolla, A., Odetti, P., Fogarty, J. & Monnier, V. M. Pentosidine formation in skin correlates with severity of complications in individuals with long-standing IDDM. Diabetes 41, 1286–1292 (1992).

5. Nagaraj, R. H., Shipanova, I. N. & Faust, F. M. Protein cross-linking by the Maillard reaction. Isolation, characterization, and in vivo detection of a lysine-lysine cross-link derived from methylglyoxal. Journal of Biological Chemistry 271, 19338–19345 (1996).

6. Martin, M. S., Jacob-Dolan, J. W., Pham, V. T. T., Sjoblom, N. M. & Scheck, R. A. The chemical language of protein glycation. Nat Chem Biol (2024) doi:10.1038/S41589-024-01644-Y.

7. Schmidt, A. M. et al. Rage: A Novel Cellular Receptor for Advanced Glycation End Products. Diabetes 45, S77–S80 (1996).

8. Laughlin, T. et al. Autophagy activators stimulate the removal of advanced glycation end products in human keratinocytes. Journal of the European Academy of Dermatology and Venereology 34, 12–18 (2020).

9. Zawada, A. et al. Accumulation of Advanced Glycation End-Products in the Body and Dietary Habits. Nutrients 14, 3982 (2022).

10. Delpierre, G. et al. Identification, cloning, and heterologous expression of a mammalian fructosamine-3-kinase. Diabetes 49, 1627–1634 (2000).

11. Collard, F., Delpierre, G., Stroobant, V., Matthijs, G. & Van Schaftingen, E. A Mammalian Protein Homologous to Fructosamine-3-Kinase Is a Ketosamine-3-Kinase Acting on Psicosamines and Ribulosamines but not on Fructosamines. Diabetes 52, 2888–2895 (2003).

12. Fortpied, J., Gemayel, R., Stroobant, V. & Van Schaftingen, E. Plant ribulosamine/erythrulosamine 3-kinase, a putative protein-repair enzyme. Biochem J 388, 795–802 (2005).

13. Fortpied, J., Gemayel, R., Stroobant, V. & Van Schaftingen, E. Plant ribulosamine/erythrulosamine 3-kinase, a putative protein-repair enzyme. Biochemical Journal 388, 795–802 (2005).

14. Gemayel, R. et al. Many fructosamine 3-kinase homologues in bacteria are ribulosamine/erythrulosamine 3-kinases potentially involved in protein deglycation. FEBS Journal 274, 4360–4374 (2007).

15. Wu, S., Lu, H. & Bai, Y. Nrf2 in cancers: A double-edged sword. Cancer Med 8, 2252 (2019).

16. Huang, Y., Li, W., Su, Z. yuan & Kong, A. N. T. The complexity of the Nrf2 pathway: beyond the antioxidant response. J Nutr Biochem 26, 1401–1413 (2015).

17. Sanghvi, V. R. et al. The Oncogenic Action of NRF2 Depends on De-glycation by Fructosamine-3-Kinase. Cell 178, 807-819.e21 (2019).

18. Garg, A. et al. The molecular basis of Human FN3K mediated phosphorylation of glycated substrates. Nature Communications 2025 16:1 16, 1–14 (2025).

19. Lokhandwala, J. et al. Structural basis for FN3K-mediated protein deglycation. Structure 32, 1711-1724.e5 (2024).

20. Harms, M. J. & Thornton, J. W. Analyzing protein structure and function using ancestral gene reconstruction. Current Opinion in Structural Biology vol. 20 360–366 Preprint at 10.1016/j.sbi.2010.03.005 (2010).

21. Thornton, J. W. Resurrecting ancient genes: Experimental analysis of extinct molecules. Nat Rev Genet 5, 366–375 (2004).

22. Trudeau, D. L., Kaltenbach, M. & Tawfik, D. S. On the Potential Origins of the High Stability of Reconstructed Ancestral Proteins. Mol Biol Evol 33, 2633–2641 (2016).

23. Taylor, S. S. & Kornev, A. P. Protein kinases: evolution of dynamic regulatory proteins. Trends Biochem Sci 36, 65–77 (2011).

24. Kornev, A. P. & Taylor, S. S. Dynamics driven allostery in protein kinases. Trends Biochem Sci 40, 628 (2015).

25. Hayden, A. et al. The Nrf2 transcription factor contributes to resistance to cisplatin in bladder cancer. Urologic Oncology: Seminars and Original Investigations 32, 806–814 (2014).

26. Shim, G. seong, Manandhar, S., Shin, D. ha, Kim, T. H. & Kwak, M. K. Acquisition of doxorubicin resistance in ovarian carcinoma cells accompanies activation of the NRF2 pathway. Free Radic Biol Med 47, 1619–1631 (2009).

27. Niture, S. K. & Jaiswal, A. K. Nrf2-induced antiapoptotic Bcl-xL protein enhances cell survival and drug resistance. Free Radic Biol Med 57, 119–131 (2013).

28. Wu, J., Zhang, L., Li, H., Wu, S. & Liu, Z. Nrf2 induced cisplatin resistance in ovarian cancer by promoting CD99 expression. Biochem Biophys Res Commun 518, 698–705 (2019).

29. Dinkova-Kostova, A. T. & Copple, I. M. Advances and challenges in therapeutic targeting of NRF2. Trends Pharmacol Sci 44, 137–149 (2023).

30. Buda, K., Miton, C. M., Fan, X. C. & Tokuriki, N. Molecular determinants of protein evolvability. Trends Biochem Sci 48, 751–760 (2023).

31. Bloom, J. D., Labthavikul, S. T., Otey, C. R. & Arnold, F. H. Protein stability promotes evolvability. Proc Natl Acad Sci U S A 103, 5869–5874 (2006).

32. Tokuriki, N., Stricher, F., Schymkowitz, J., Serrano, L. & Tawfik, D. S. The Stability Effects of Protein Mutations Appear to be Universally Distributed. J Mol Biol 369, 1318–1332 (2007).

33. Tokuriki, N., Stricher, F., Serrano, L. & Tawfik, D. S. How Protein Stability and New Functions Trade Off. PLoS Comput Biol 4, e1000002 (2008).

34. Mashiba, S., Uchida, K., Okuda, S. & Tomita, S. Measurement of glycated albumin by the nitroblue tetrazolium colorimetric method. Clinica Chimica Acta 212, 3–15 (1992).

35. Walker, S. W., Howie, A. F. & Smith, A. F. The measurement of glycosylated albumin by reduction of alkaline nitro-blue tetrazolium. Clinica Chimica Acta 156, 197–206 (1986).

36. Consortium, T. U. et al. UniProt: the Universal Protein Knowledgebase in 2025. Nucleic Acids Res 53, D609–D617 (2025).

37. Potter, S. C. et al. HMMER web server: 2018 update. Nucleic Acids Res 46, W200–W204 (2018).

38. Li, W. et al. The EMBL-EBI bioinformatics web and programmatic tools framework. Nucleic Acids Res 43, W580–W584 (2015).

39. Madeira, F. et al. The EMBL-EBI Job Dispatcher sequence analysis tools framework in 2024. Nucleic Acids Res 52, W521–W525 (2024).

40. Guindon, S. et al. New Algorithms and Methods to Estimate Maximum-Likelihood Phylogenies: Assessing the Performance of PhyML 3.0. Syst Biol 59, 307–321 (2010).

41. Lefort, V., Longueville, J. E. & Gascuel, O. SMS: Smart Model Selection in PhyML. Mol Biol Evol 34, 2422–2424 (2017).

42. Yang, Z. PAML 4: Phylogenetic Analysis by Maximum Likelihood. Mol Biol Evol 24, 1586–1591 (2007).

43. Sievers, F. et al. Fast, scalable generation of high-quality protein multiple sequence alignments using Clustal Omega. Mol Syst Biol 7, (2011).

44. Crooks, G. E., Hon, G., Chandonia, J. M. & Brenner, S. E. WebLogo: A sequence logo generator. Genome Res 14, 1188–1190 (2004).

45. Abramson, J. et al. Accurate structure prediction of biomolecular interactions with AlphaFold 3. Nature 2024 630:8016 630, 493–500 (2024).

46. Felline, A., Seeber, M. & Fanelli, F. webPSN v2.0: a webserver to infer fingerprints of structural communication in biomacromolecules. Nucleic Acids Res 48, W94–W103 (2020).

47. Ferruz, N., Schmidt, S. & Höcker, B. ProteinTools: a toolkit to analyze protein structures. Nucleic Acids Res 49, W559–W566 (2021).

