## Supplemental Figures for "Ancestral protein reconstruction reveals the mechanism of substrate specificity in FN3K-mediated deglycation"

**Figure S1.** Kinase activity of substrate binding mutations in HsFN3K

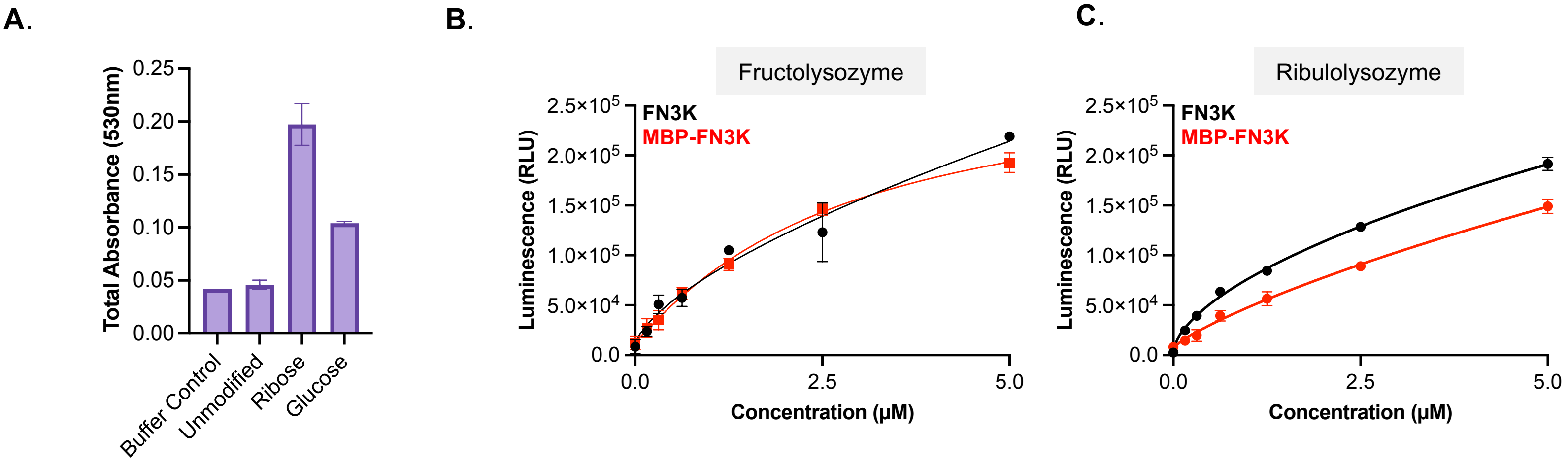


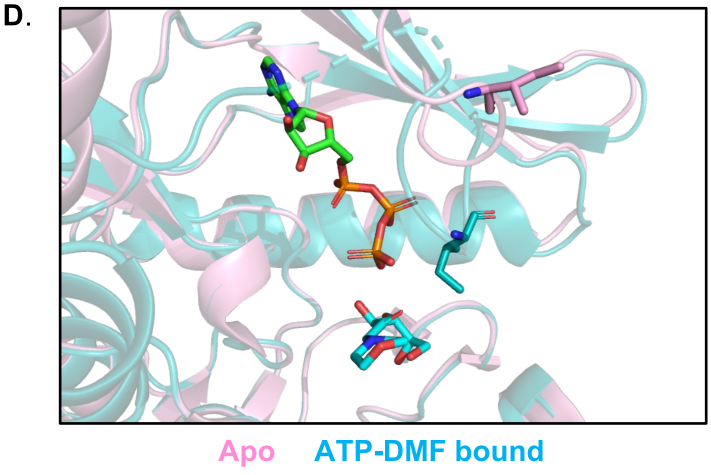


E.

| Protein | +ATP (Δ˚C) |
|---|---|
| FN3K | +4.2 |
| I25A | +0.0 |
| D217A | +2.3 |
| W219A | +5.1 |
| F252A | +2.3 |
| N287A | +0.8 |
| H288A | +4.2 |
| H291A | +3.1 |
| F292A | +1.5 |

**Figure S2**. Ancestral reconstruction of fructosamine kinase family generates stable proteins with high homogeneity.

**A.**

```
HsFN3K       --------MEQLLRAELRTATLRAFGGPGAGCISEGRAYDTDAGPVFVKVNRRTQARQMF  52
HsKT3K       --------MEELLRRELGCSSVRATGHSGGGCISQGRSYDTDQGRVFVKVNPKAEARRMF  52
ancFN3K      MAAMSEDPMEALLKRELGTAVLKATGHSGGGCISQGQSYDTDRGRVFVKINHKAEARRMF  60
ancKT3K2     MAAMSEDPMEALLKRELGTAVLKATGHSGGGCISQGQSYDTDRGRVFVKINSKAEARRMF  60
ancKT3K1     MAAMSEDPMEALLKRELGTAVLKATGHSGGGCISQGQSYDTDRGRVFVKINHKAEARRMF  60
ancFN3KRP    MAAMSEDPMEALLKRELGTAVLKATGHSGGGCISQGQSYDTDRGRVFVKINHKAEARRMF  60

HsFN3K       EGEVASLEALRSTGLVRVPRPMKVIDLPGGGAAFVMEHLKMKSLSSQASKLGEQMADLHL  112
HsKT3K       EGEMASLTAILKTNTVKVPKPIKVLDAPGGGSVLVMEHMDMRHLSSHAAKLGAQLADLHL  112
ancFN3K      EGEMASLEAILKTNTVKVPKPIKVIDLPGGGAVFVMEHLDMRGLSKHSAKLGEQLADLHL  120
ancKT3K2     EGEMASLEAILKTETVKVPKPIKVIDLPGGGAVLVMEHLDMRGLSRHSAKLGTQLADLHL  120
ancKT3K1     EGEMASLEAILKTETVKVPKPIKVIDLPGGGAVLVMEHLDMRGLSRHSAKLGEQLADLHL  120
ancFN3KRP    EGEMASLEAILKTETVKVPKPIKVIDLPGGGAVFVMEHLDMRGLSKHSAKLGEQLADLHL  120

HsFN3K       YNQKLREKLKEEENTVGRRGEGAEPQYVDKFGFHTVTCCGFIPQVNEWQDDWPTFFARHR  172
HsKT3K       DNKKLGEMRLKEAGTVGRGGGQEERPFVARFGFDVVTCCGYLPQVNDWQEDWVVFYARQR  172
ancFN3K      HNQKLGEKLKKEAGTVGKGAGQSEPQYVDKFGFHTVTCCGYIPQVNEWQDDWPTFFARHR  180
ancKT3K2     HNQKLGEKLKKEAGTVGKGAGQSEVQFVDQFGFHTVTCCGYLPQVNDWQDDWVTFFARQR  180
ancKT3K1     HNQKLGEKLKKEAGTVGKGAGQSEVQYVDKFGFHTVTCCGYLPQVNDWQDDWVTFFARQR  180
ancFN3KRP    HNQKLGEKLKKEAGTVGKGAGQSEVQYVDKFGFHTVTCCGYLPQVNEWQDDWVTFFARQR  180

HsFN3K       LQAQLDLIEKDYADREARELWSRLQVKIPDLFCGLEIVPALLHGDLWSGNVAEDDV-GPI  231
HsKT3K       IQPQMDMVEKESGDREALQLWSALQLKIPDLFRDLEIIPALLHGDLWGGNVAEDSS-GPV  231
ancFN3K      LQAQLDLIEKDYGDREARELWSQLQLKIPDLFCDVEIVPALLHGDLWGGNVAEDDSGEPI  240
ancKT3K2     IQPQMDMIEKRSGDREARELWSQLQLKIPDLFCDMEIVPALLHGDLWGGNVAEDDSGEPI  240
ancKT3K1     IQPQMDMIEKRYGDREARELWSQLQLKIPDLFCDMEIVPALLHGDLWGGNVAEDDSGEPI  240
ancFN3KRP    LQPQLDMIEKRYGDREARELWSQLQLKIPDLFCDVEIVPALLHGDLWGGNVAEDDSGEPI  240

HsFN3K       IYDPASFYGHSEFELAIALMFGGFPRSFFTAYHRKIPKAPGFDQRLLLYQLFNYLNHWNH  291
HsKT3K       IFDPASFYGHSEYELAIAGMFGGFSSSFYSAYHGKIPKAPGFEKRLQLYQLFHYLNHWNH  291
ancFN3K      IFDPASFYGHSEFELAIAGMFGGFSSSFFSAYHSKIPKAPGFEKRLKLYQLFNYLNHWNH  300
ancKT3K2     IFDPASFYGHSEYELAIAGMFGGFSSSFYSAYHSKIPKAPGFEKRLKLYQLFHYLNHWNH  300
ancKT3K1     IFDPASFYGHSEYELAIAGMFGGFSSSFYSAYHSKIPKAPGFEKRLKLYQLFHYLNHWNH  300
ancFN3KRP    IFDPASFYGHSEYELAIAGMFGGFSSSFYSAYHSKIPKAPGFEKRLKLYQLFHYLNHWNH  300

HsFN3K       FGREYRSPSLGTMRRLLK----  309
HsKT3K       FGSGYRGSSLNIMRNLVK----  309
ancFN3K      FGTGYRSSSLNIMRKLLKCLKA  322
ancKT3K2     FGTGYRGSSLNIMRNLVKCLKA  322
ancKT3K1     FGTGYRSSSLNIMRNLLKCLKA  322
ancFN3KRP    FGTGYRSSSLNIMRNLLKCLKA  322
```

**B.**

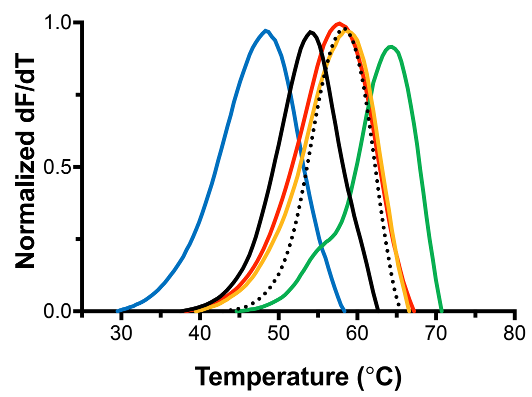


**C.**

| | |
|---|---|
| HsFN3K | 54.4˚C |
| HsKT3K | 58.4˚C |
| ancFN3KRP | 57.7˚C |
| ancKT3K$^1$ | 58.7˚C |
| ancKT3K$^2$ | 64.8˚C |
| ancFN3K | 47.9˚C |

**Figure S3.** Switch mutations in ancFN3Ks.

**A.**

| Site | ancFN3KRP | ancFN3K | KT3K | FN3K | Site |
|---|---|---|---|---|---|
| 74 | E | N | N | G | 66 |
| 145 | V | P | R | P | 137 |
| 162 | L | I | L | I | 154 |
| 173 | V | P | V | P | 165 |
| 179 | Q | H | Q | H | 171 |
| 183 | P | A | P | A | 175 |
| 187 | M | L | M | L | 179 |
| 191 | R | D | E | D | 183 |
| 253 | Y | F | Y | F | 244 |
| 269 | Y | F | Y | F | 260 |
| 293 | H | N | H | N | 284 |
| 315 | N | K | N | R | 306 |

**B.**

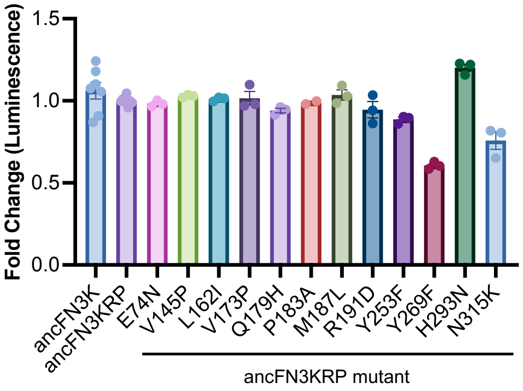


**C.**

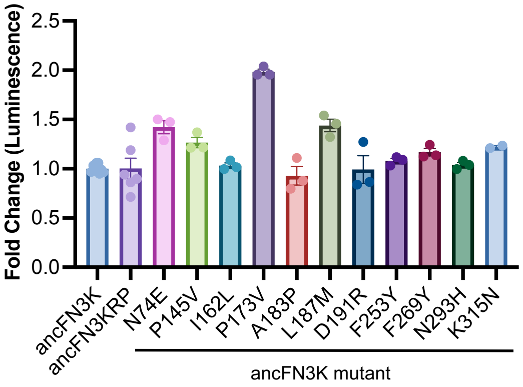

**Figure S4.** Mutations in FN3K make them more KT3K-like.

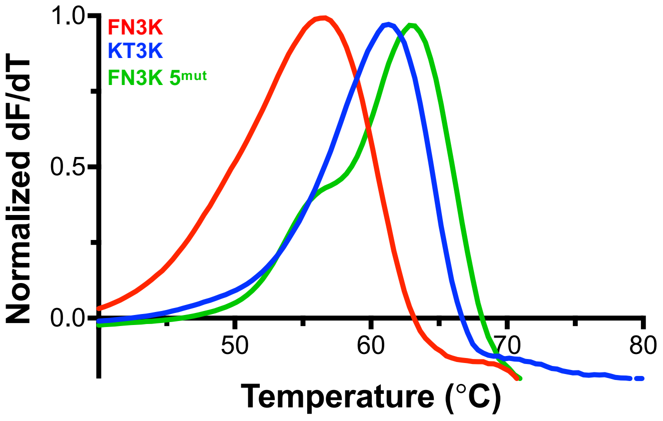

**Figure S5.** Intramolecular contact analysis of HsFN3K and HsKT3K

**A.**

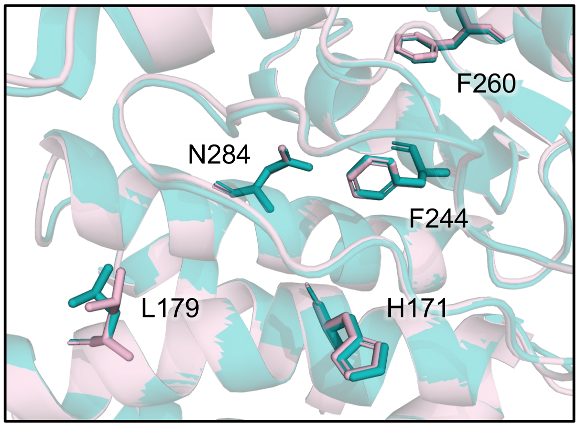


**B.**

| HsFN3K N284 | |
|---|---|
| F283 | 1.33 |
| Y285 | 1.34 |
| **H288** | **2.91** |
| Q281 | 2.97 |
| Y280 | 3.08 |
| L282 | 3.19 |
| **N287** | **3.30** |
| L286 | 3.33 |
| M251 | 3.49 |
| I248 | 3.64 |
| A247 | 4.06 |
| **F244** | **4.29** |
| **F252** | **4.79** |
| W289 | 4.99 |
| **D217** | **5.16** |
| L279 | 5.73 |
| Y296 | 5.86 |

| HsKT3K H284 | |
|---|---|
| F283 | 1.33 |
| Y285 | 1.34 |
| **H288** | **2.83** |
| Q281 | 3.13 |
| Y280 | 3.30 |
| L282 | 3.31 |
| **Y244** | **3.32** |
| L286 | 3.41 |
| **N287** | **3.43** |
| I248 | 3.53 |
| M251 | 3.79 |
| G216 | 4.03 |
| A247 | 4.14 |
| C151 | 4.71 |
| **D217** | **4.75** |
| W289 | 5.01 |
| **F252** | **5.20** |
| Y168 | 5.82 |
| Y296 | 5.93 |
| E245 | 5.93 |

**Figure S6.** Protein Structure Network analysis of HsFN3K and AtFN3K

**A.** AtFN3K HsFN3K Shared

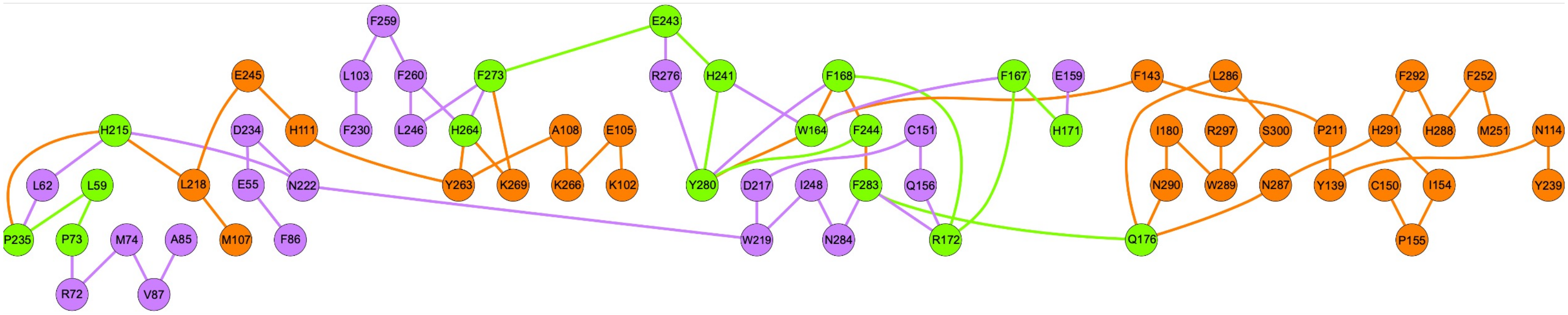


**B.**

```
AtFN3K      MAVASLSICFSARPHLLLRNFSPRPKFVAMAAMSEDPIREWILTEGKATQITKIGSVGGG    60
HsFN3K      -------------------------------------MEQLLRAELRTATLRAFGGPGAG    23
                                                 :.: : :* ::: :  :*. *.*

AtFN3K      CINLASHYQTDAGSFFVKTNRS-IGPAMFEGEALGLEAMYETRTIRVPNPHKAGELPTGG    119
HsFN3K      CISEGRAYDTDAGPVFVKVNRRTQARQMFEGEVASLEALRSTGLVRVPRPMKVIDLPGGG    83
            **. .  *:**** .***.**   .  *****. .***: .*  :***.* *. :** **

AtFN3K      SYIIMEFIDFGGSRGNQAELGRKLAEMHKAG-----------------------KTSKGF    156
HsFN3K      AAFVMEHLKMKSLSSQASKLGEQMADLHLYNQKLREKLKEEENTVGRRGEGAEPQYVDKF    143
            : ::**.:.: .  .: ::**.::*::*  .                       :  . *

AtFN3K      GFEVDNTIGSTPQINTWSSDWIEFYGEKRLGYQLKLARDQYGDSAIYQKGHTLIQNMAPL    216
HsFN3K      GFHTVTCCGFIPQVNEWQDDWPTFFARHRLQAQLDLIEKDYADREARELWSRLQVKIPDL    203
            **.. .  *  **:* *..**  *:..:**  **.* ..:*.*    :    *  ::  *

AtFN3K      FENVVIEPCLLHGDLWSGNIAYDKNNEPVILDPACYYGHNEADFGMSW-CAGFGESFYNA    275
HsFN3K      FCGLEIVPALLHGDLWSGNVAED-DVGPIIYDPASFYGHSEFELAIALMFGGFPRSFFTA    262
            * .: * *.**********:* * :  *:* ***.:***.* ::.::   .** .**:.*

AtFN3K      YFKVMPKQAGYEKRRDLYLLYHYLNHYNLFGSGYRSSAMSIIDDYLRMLKA      326
HsFN3K      YHRKIPKAPGFDQRLLLYQLFNYLNHWNHFGREYRSPSLGTMRRLLK----      309
            *.: :**  *:::*  ** *::****:* **  *** ::. :   *:
```

**Figure S7.** Neofunctionalization of fructosamine repair by FN3Ks begins with a destabilization event.

**A.**

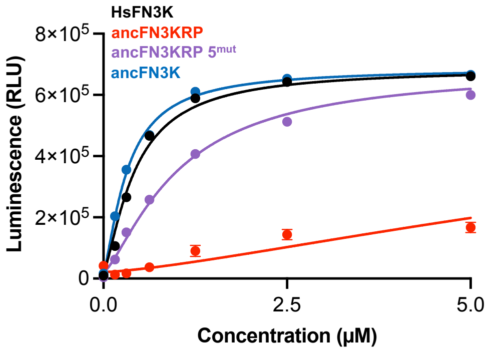


**B.**

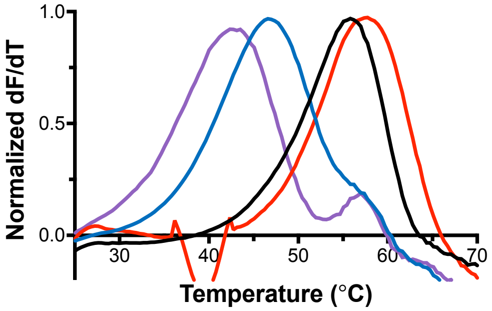


**C.**

| | |
|---|---|
| HsFN3K | 56.2˚C |
| ancFN3KRP | 57.7˚C |
| ancFN3KRP 5mut | 41.5˚C |
| ancFN3K | 46.2˚C |
