## Supplemental Figure Legends for "Ancestral protein reconstruction reveals the mechanism of substrate specificity in FN3K-mediated deglycation"

**Figure S1. Kinase activity of substrate binding mutations in HsFN3K. A.** NBT colorimetric assay of buffer control, unmodified lysozyme, ribose-glycated lysozyme, and glucose-glycated lysozyme. **B-C.** Kinase assay of MBP-tagged and untagged FN3K on fructolysozyme (B) and ribulolysozyme (C). Each data point represents means of triplicates; error bars indicate standard error. **D.** HsFN3K I25 position in apo (PDB: 9CX8, pink) and ATP-DMF bound (PDB: 9CXM, blue). **E.** Change in melting temperatures of FN3K mutants in the presence of ATP.

**Figure S2. Ancestral reconstruction of fructosamine kinase family generates stable proteins with high homogeneity. A.** Multiple sequence alignment of HsFN3K, HsKT3K, ancFN3KRP, ancKT3K, ancKT3K, and ancFN3K. Conserved amino acid residues across all proteins are highlighted in gray. Amino acid residues found only in KT3K-like proteins are highlighted in yellow. Substrate binding residues from Figure 1are indicated by black arrows. **B.** Normalized DSF melt curves of HsFN3K (black solid), HsKT3K (dotted), ancFN3KRP (red), ancKT3K$^{1}$ (yellow), ancKT3K$^{2}$ (green), and ancFN3K (blue). **C.** Melting temperatures determined in (B).

**Figure S3. Switch mutations in ancFN3Ks. A.** The 12 amino acid residues that differ between ancFN3KRP and ancFN3KRP and their corresponding residues in HsFN3K and HsKT3K. **B-C.** Kinase activity of ancFN3KRP mutants (B) and ancFN3K mutants (C) towards ribulolysozyme. Each bar graph represents the activity at 5 µM of enzyme. Each data point represents means of triplicates; error bars indicate standard error.

**Figure S4. Mutations in FN3K make them more KT3K-like. A.** DSF melt curve of HsFN3K, HsKT3K, and HsFN3K $5^{mut}$.

**Figure S5. Intramolecular contact analysis of HsFN3K and HsKT3K. A.** HsFN3K apo (pink, PDB: 9CX8) versus ATP-DMF bound (blue, PDB: 9CXM) structures. H171, L179, F244, F260, and N284 are represented as sticks. **B.** Intramolecular contact analysis of HsFN3K N284 (B) and AlphaFold predicted HsKT3K H284 (C). Purple represents established substrate-binding residues. Pink represents residues highlighted in this study.

**Figure S6. Protein Structure Network analysis of HsFN3K and AtFN3K. A.** Difference PSN of AtFN3K (PDB: 6OID, purple) and HsFN3K (PDB: 9CX8, orange). Shared residues in each network are green. Residues are labeled as HsFN3K residues. **B.** Multiple sequence alignment of HsFN3K and AtFN3K. Yellow box highlights alpha-helix where F244 resides.

**Figure S7. Neofunctionalization of fructosamine repair by FN3Ks begins with a destabilization event. A.** Kinase assay of ancFN3KRP, ancFN3KRP $5^{mut}$, ancFN3K, and HsFN3K against DMF. Each data point represents means of triplicates; error bars indicate standard error. **B**. DSF melting curves and corresponding melting temperatures for ancFN3KRP, ancFN3KRP $5^{mut}$, ancFN3K, and HsFN3K.
